# A non-canonical chemical feedback self-limits nitric oxide-cyclic GMP signaling in health and disease

**DOI:** 10.1101/383208

**Authors:** Vu Thao-Vi Dao, Mahmoud H. Elbatreek, Martin Deile, Pavel I. Nedvetsky, Andreas Güldner, César Ibarra-Alvarado, Axel Gödecke, Harald H.H.W. Schmidt

**Affiliations:** Department of Pharmacology and Personalised Medicine, MeHNS, FHML, Maastricht University, Maastricht, The Netherlands; Department of Pharmacology, Johannes Gutenberg University, Mainz, Germany; Department for Pharmacology and Toxicology, Faculty of Pharmacy, Zagazig University, Zagazig, Egypt; Asklepios-Klinik Radeberg, Dresden, Germany; Universitätsklinikum Münster, Medical Cell Biology, Münster, Germany; Residency Anaesthesiology, Department of Anaesthesiology und Critical Care Medicine, Technische Universität, Dresden, Germany; Facultad de Química, Universidad Autónoma de Querétaro, Mexico; Institut für Herz- und Kreislaufphysiologie Heinrich-Heine-Universität, Düsseldorf, Germany

**Keywords:** endothelial cells, soluble guanylate cyclase, nitric oxide, cyclic guanosine mono phosphate, redox regulation, Bay 58-2667, apo-sGC

## Abstract

Nitric oxide (NO)-cyclic GMP (cGMP) signaling is a vasoprotective pathway therapeutically targeted for example in pulmonary hypertension. Its dysregulation in disease is incompletely understood. Here we show in pulmonary artery endothelial cells that feedback inhibition by NO of the NO receptor, the cGMP forming soluble guanylate cyclase (sGC), may contribute to this. Both endogenous NO from endothelial NO synthase or exogenous NO from NO donor compounds decreased sGC protein and activity. This was not mediated by cGMP as the NO-independent sGC stimulator or direct activation of cGMP-dependent protein kinase did not mimic it. Thiol-sensitive mechanisms were also not involved as the thiol-reducing agent, N-acetyl-L-cysteine did not prevent this feedback. Instead, both *in-vitro* and *in-vivo* and in health and acute respiratory lung disease, chronically elevated NO led to the inactivation and degradation of sGC whilst leaving the heme-free isoform, apo-sGC, intact or even increasing its levels. Thus, NO regulates sGC in a bimodal manner, acutely stimulating and chronically inhibiting, as part of self-limiting direct feedback that is cGMP-independent. In high NO disease conditions, this is aggravated but can be functionally recovered in a mechanism-based manner by apo-sGC activators that re-establish cGMP formation.

## Introduction

The nitric oxide (NO)-cGMP signaling pathway plays several important roles in physiology including cardiopulmonary homeostasis ^1,2^. The main receptor and mediator of NO’s actions is soluble guanylate cyclase (sGC), a heterodimeric heme protein. In its Fe(II)heme-containing state, sGC binds NO and is thereby activated to convert guanosine triphosphate (GTP) to the second messenger, cGMP, whose steady state levels are counter-regulated by different phosphodiesterases (PDEs) ^3^. cGMP exerts its cardiopulmonary effects via activating cGMP-dependent protein kinase-I (PKG) ^4^. This results in potent vasodilatory, anti-proliferative and anti-thrombotic effects ^5^. In disease, heme loss, appearance of NO-insensitive apo-sGC and impaired NO-cGMP signaling have been described ^6,7^.

In addition to sGC’s acute activation, NO appears to have further roles in regulating sGC. During enzyme maturation, NO facilitates heme incorporation into sGC ^8,9^, and activation of sGC by NO is followed by an acute and rapid desensitization involving protein S-nitrosylation ^10,11^. In addition, chronic exposure to NO donor drugs has been suggested to negatively affect sGC activity in a not fully reversible manner ^12-14^. It is unclear, however, whether this effect pertains also to endogenously formed NO and has pathophysiological relevance.

Here, we examine this important knowledge gap in the (patho)biology of NO. As model systems we chose porcine pulmonary artery endothelial cells (PPAECs) as they relate to the clinical application of NO and cGMP modulating drugs in pulmonary hypertension ^15,16^. We investigate the effects of chronic exposure to both exogenous (from NO donor drugs) and endogenous NO on sGC protein and activity. In addition, we investigate in health and disease whether chronic effects of NO on sGC involve canonical cGMP signaling, thiol modulation, or formation of heme-free, apo-sGC. As disease model, we use again a condition related to pulmonary hypertension and chronically elevated levels of NO, porcine acute respiratory disease syndrome (ARDS) ^17–19^.

## Results

### NO chronically decreases vascular sGC protein and activity *in-vivo* and *in-vitro*

To analyze the chronic effects of NO at a mechanistic level, we studied PPAECs. Cells were incubated for up to 72h in the presence of the NO synthase (NOS) inhibitor N^G^-nitro L-arginine methyl ester (L-NAME) and sGC expression and activity were measured. In the presence of L-NAME to eliminate endogenous NO formation, protein levels of the heme-binding sGCβ_1_ subunit were increased (Fig. 1A). This was associated also with increased sGC activity (Fig. 1B). Next we wanted to test the reverse, i.e. whether an increase of NO to supra-physiological concentrations ^20–22^ by chronic exposure to the long-acting NO donor compound, DETA/NO, would downregulate sGC. Indeed, pre-incubating cells with DETA/NO (100 μM) decreased both sGCα_1_ and sGCβ1 protein (Fig. 1C) and sGC activity (Fig. 1D). Thus, *in-vitro* in PPAECs, endogenous NO chronically downregulates sGC protein and activity in an L-NAME reversible manner, which is further aggravated by exogenous, pharmacologically applied NO in supra-physiological concentrations.

**Fig. 1.**
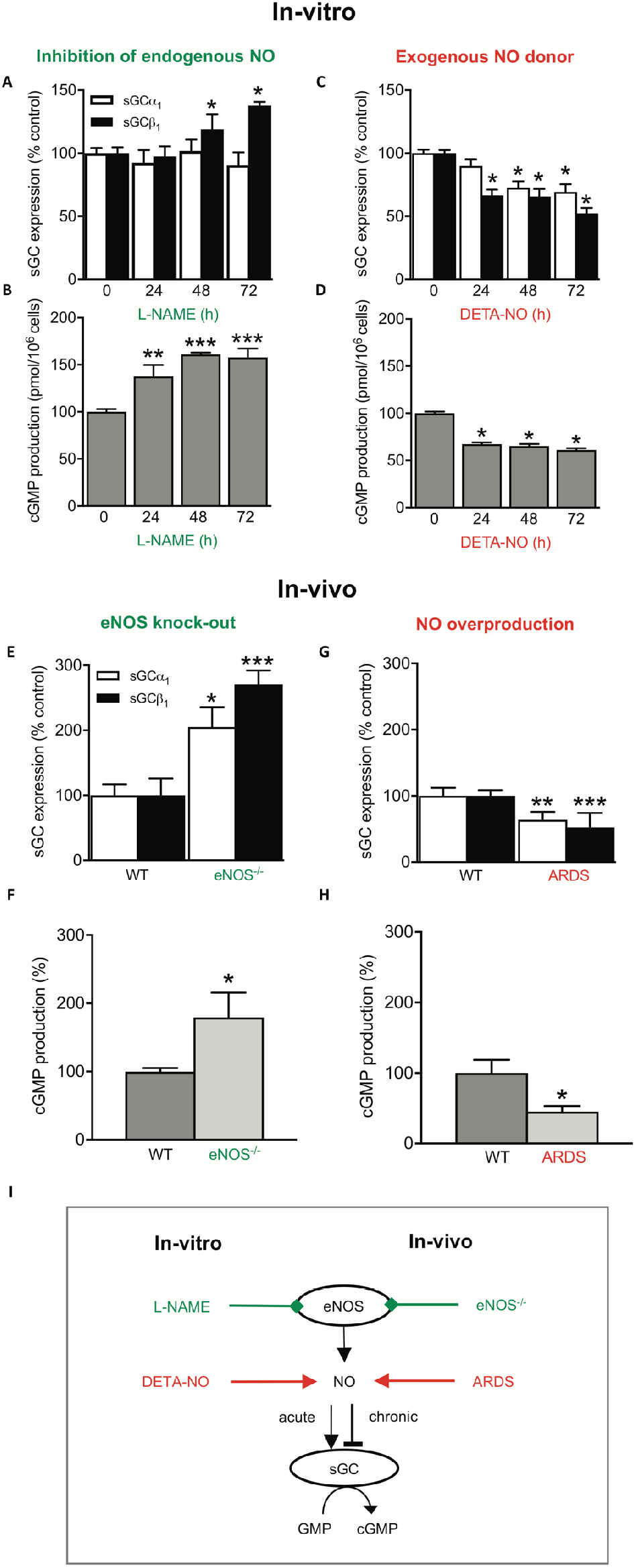
Chronic NO decreases vascular sGC protein and activity *in-vivo* and *in-vitro*. **(A)** Inhibiting basal NO formation in PPAEC by L-NAME (100 μM) for up to 72h increased sGCβ_1_ expression (N=6). **(B)** This upregulation was associated with increased sGC activity (N=3). Exposing cells to supra-physiological levels of NO by chronic exposure to the NO donor compound, DETA/NO (100 μM), for up to 72h decreased both sGCα_1_ and sGCβ_1_ protein **(C)**(N=6) and sGC activity **(D)**(N=5). *In-vivo* validation of the *in-vitro* observations showed in eNOS knock-out mice (eNOS^−/−^) mice increased sGC protein **(E)** and activity levels **(F)**(N=9), and in a porcine lung disease model (ARDS) characterized by NO overproduction, decreased sGCα_1_ and sGCβ_1_ protein **(G)**(N=5) and sGC activity levels **(H)**(N=3). Data are expressed as mean ± SEM. *, **, ***: p < 0.05, 0.01 or 0.001 *vs.* control, respectively. **(I)** Schematic summary showing that both *in-vitro* (porcine lung endothelial cells) and *in-vivo* (the porcine lung disease model, ARDS) both endogenous and exogenous NO downregulate sGC protein and activity. Representative full-length blots are presented in Supplementary Figure S3.

Next, we wanted to validate these *in-vitro* observations at an *in-vivo* level in eNOS knock-out mice (eNOS^−/−^), eliminating endogenous NO formation similar to the *in-vitro* L-NAME experiment, and in a porcine lung disease model (ARDS) characterized by NO overproduction, mimicking the exposure the supraphysiological NO concentration through the NO donor ^17,19,23^. In line with our observations in PPAECs, eNOS^−/−^ mice showed increased protein levels of sGCα_1_ and sGCβ_1_ (Fig. 1E) and increased sGC-activity (Fig. 1F), and in the high-NO pulmonary disease model, sGCα_1_ and sGCβ_1_ protein levels (Fig. 1G) and sGC activity were decreased (Fig. 1H). Collectively, these data suggest that both *in-vitro* and *in-vivo* lowering endogenous NO increases, and increasing endogenous NO lowers sGC protein subunit levels and sGC activity (Fig. 1I).

### cGMP/PKG do not mediate the downregulation of sGC protein and activity by chronic NO

Next, we aimed to clarify the mechanisms underlying the downregulation of sGC protein and activity by chronic NO. First, we tested whether cGMP/PKG signaling is involved as it had been shown previously to decrease both sGC activity ^24^ and expression ^25^. Of experimental importance, cell passaging can cause downregulation of PKG and prevent the detection of PKG-dependent signaling ^26–29^. Hence, we therefore restricted our studies to low passage number cells and ensured fully functional PKG signaling by validating the known autoregulation of PKG expression ^30,31^. Indeed, in our PPAEC system, both the PKG activator, 8-Br-cGMP, and the NO-independent sGC stimulator and PDE inhibitor, YC-1, ^32^ were able to reduce PKG expression (Supplementary Fig. S1 online) confirming the presence of fully functional PKG. We then studied whether the observed downregulation of sGC protein and activity by NO can be mimicked by cGMP or is prevented by inhibiting PKG. When we exposed PPAECs, however, for 72h with different concentrations of the sGC stimulator and PDE inhibitor, YC-1, to raise cGMP in an NO-independent manner, or to the direct PKG activator, 8-Br-cGMP, neither sGC protein nor activity were lowered (*cf.* to Fig. 1). In fact, we observed even a slight upregulation of sGC protein (Fig. 2, A and B). Consistent with this, the NO induced downregulation of sGC could not be prevented by co-incubation with the PKG inhibitor, Rp-8-Br-PET-cGMPS (Supplementary Fig. S2 online). To extend these *in-vitro* findings also to the *in-vivo* level, we subsequently studied sGC expression and activity in PKG knock-out mice (PKG^−/−^) ^33^. Consistent with our *in-vitro* data, sGC protein levels (Fig. 2D) and sGC activity (Fig. 2E) were unchanged in PKG^−/−^ as compared to wildtype mice. In conclusion, both our *in-vivo* and *in-vitro* data suggested that the downregulation of sGC protein and activity by chronic NO is cGMP- and PKG-independent and thus appeared to be due to a non-canonical mechanism (Fig. 2C). Two cGMP-independent effects on sGC have been reported, rapid desensitization ^10,20,34^, which is reversible in a thiol-dependent manner ^35,36^, and oxidative heme-loss yielding the NO-insensitive apo-form of (apo-sGC) ^7,37^. These possibilities were tested in our two next sets of experiments.

**Fig. 2.**
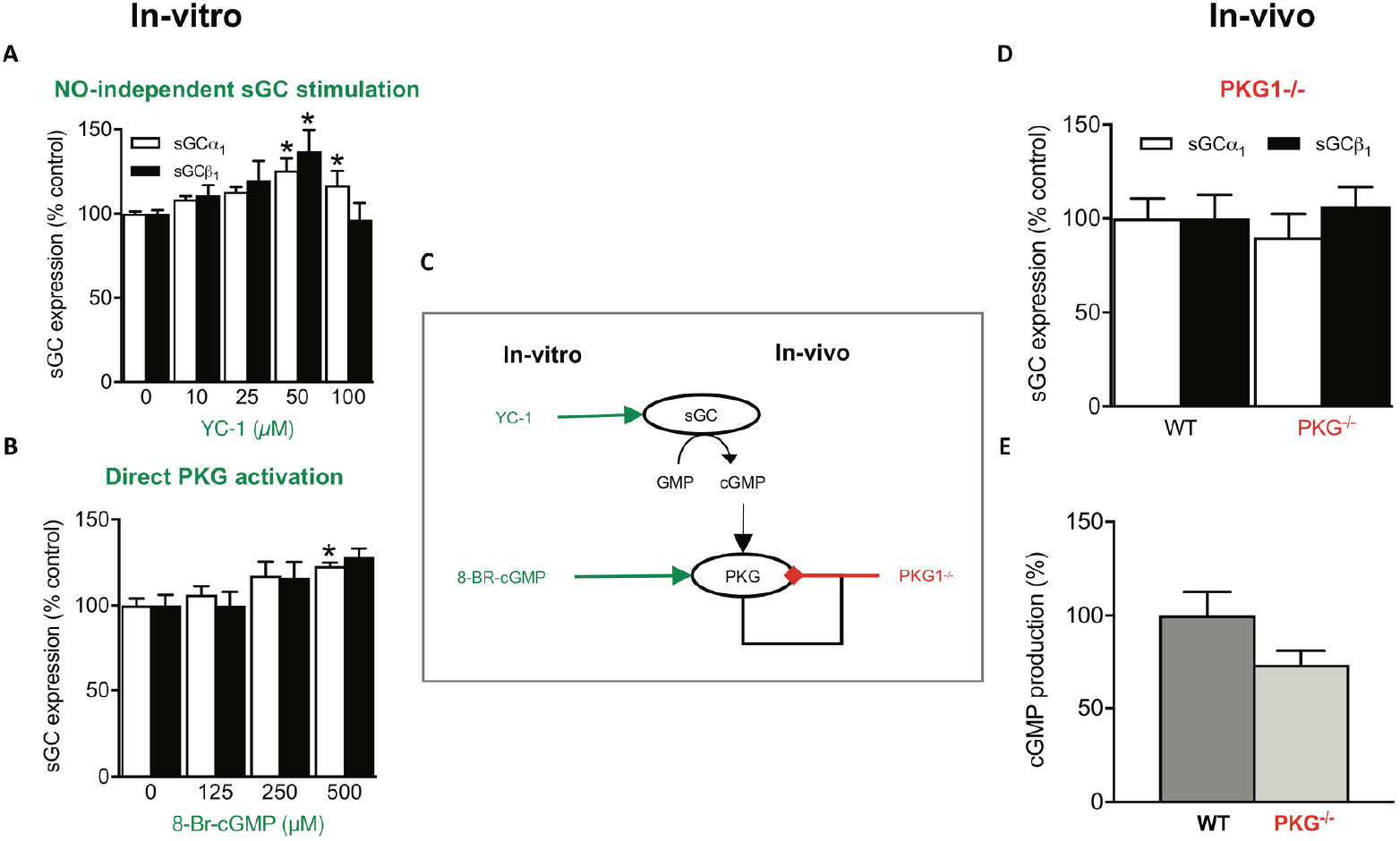
PKG does not mediate the downregulation of sGC protein and activity by chronic NO. When PPAEC were incubated for 72 h in the absence or presence of increasing concentrations of **(A)** the NO-independent sGC stimulator, YC-1 (N=6), this did not cause a downregulation of sGCα_1_ and sGCβ_1_ expression but rather a small upregulation. Consistent with this, in **(B)** the direct PKG activator 8-Br-cGMP (N=6) lead to increased sGCα_1_ protein expression. **(C)** The scheme summarizes the *in-vivo* and *in-vitro* data suggesting that the downregulation of sGC protein and activity by chronic NO is cGMP- and PKG-independent and thus appeared to be due to a non-canonical mechanism. **(D)** sGC protein expression (N=4) and **(E)** activity (N=4) are not altered in PKG^−/−^ as compared to wildtype mice. Data are expressed as mean ± SEM. *,**,***: p < 0.05, 0.01 or 0.001 *vs.* control, respectively. Representative full-length blots are presented in Supplementary Figure S3.

### NO-induced sGC downregulation is thiol-independent

It has been shown previously, that thiol-sensitive mechanisms are involved in sGC regulation such as sGC maturation and airway pathologies such as asthma ^20^. Therefore, we assessed whether NO-posttranslational modification of free-thiol cysteines i.e. S-nitrosylation contributes to the downregulation of sGC by high chronic NO incubation. For this approach, PPAECs were again exposed for 72 h to DETA-NO (100μM) in absence or presence, over the full-time frame, of the membrane-permeable thiol-reducing agent, N-acetyl-L-cysteine (NAC; 1mM). NAC is a membrane-permeable de-nitrosylating agent and glutathione precursor that has been shown to protect from sGC nitrosylation ^38,39^. Although some studies had used higher concentration of NAC, 1 mM is sufficient to induce de-nitrosylation ^40,41^. The presence of NAC, however, neither affected sGC protein levels (Fig. 3A) nor sGC activity (Fig. 3B). This set of experiments suggested that it is unlikely that a thiol-reversible mechanism similar to the acute desensitization is involved in the chronic NO-induced downregulation of sGC protein and activity. This left oxidative heme-loss yielding the NO-insensitive apo-form of (apo-sGC) ^7,37^ as only known cGMP-independent effect on sGC.

**Fig. 3.**
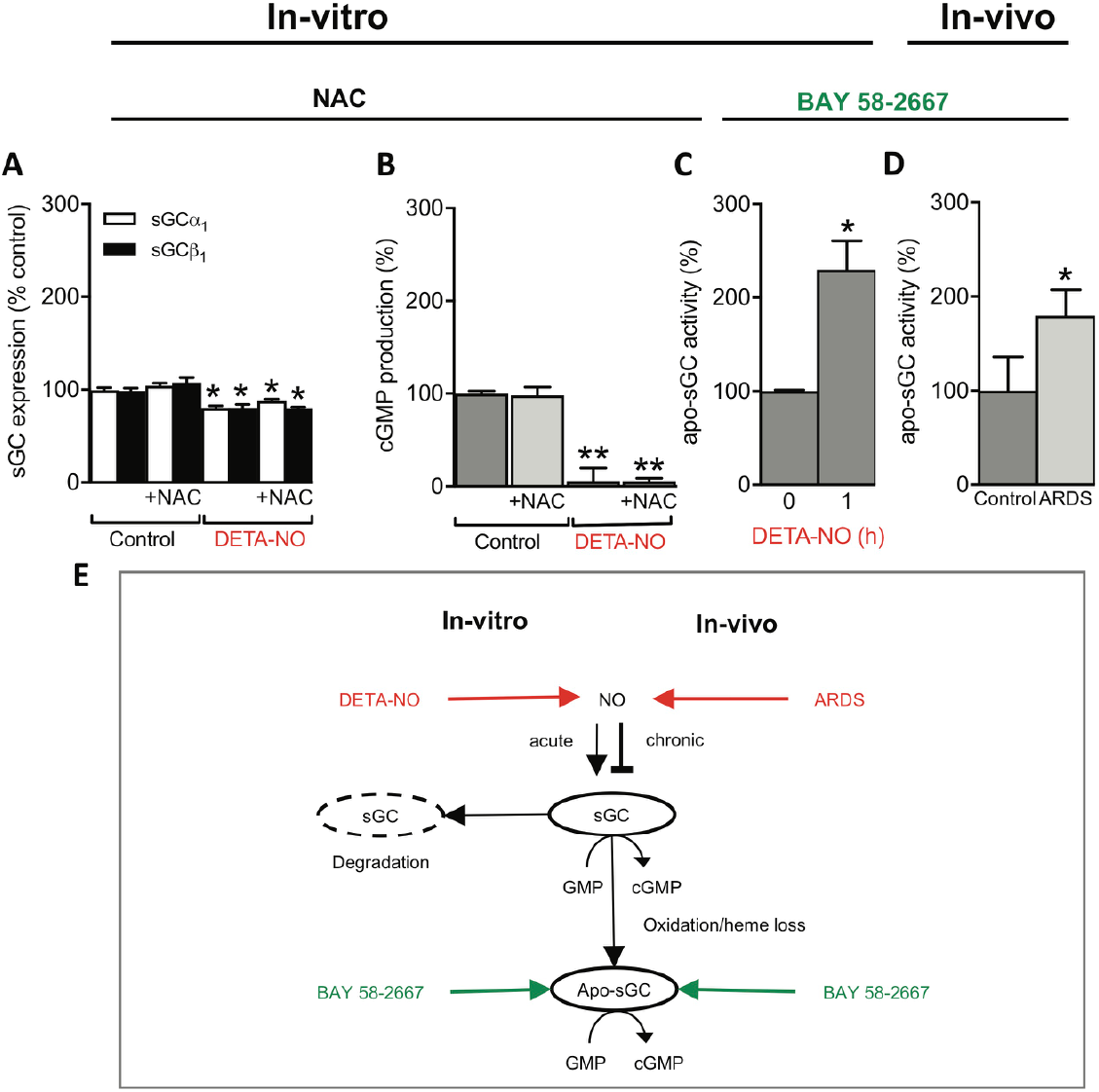
NO-induced sGC downregulation is thiol-independent but involves sGC loss and a shift towards apo-sGC. When PPAECs were exposed for 72 h to DETA-NO (100μM) in the absence and presence of N-acetyl-L-cysteine (NAC; 1mM), NAC neither affected sGC protein levels (N=5) (**A**) nor activity (N=4) (**B**). Exposure of PPAECs for 72h to DETA-NO (100 μM) increased apo-sGC activity, measured as BAY 58-2667-induced cGMP formation (BAY 58-2667, 10 μM) (N=3) (**C**). Validation of the above *in-vitro* mechanistic findings *in-vivo* in the porcine high-NO ARDS model showing also increased apo-sGC activity (N=3) **(D)**. **(E)** A scheme summarizing both our *in-vitro* and *in-vivo* data that both endogenous NO or pharmacological NO donor compounds that acutely stimulate sGC, chronically decreased both sGC protein and activity leading to inactivation of sGC and an apparent net shift towards NO-insensitive apo-sGC. Data are expressed as mean ± SEM. *,**: p < 0.05 or 0.01 *vs.* control, respectively. Representative full-length blots are presented in Supplementary Figure S3.

### NO-induced sGC downregulation generates NO-insensitive sGC

We therefore examined whether chronic NO converts sGC to the NO-insensitive, heme-free form of sGC, i.e. apo-sGC. To assay for the presence of apo-sGC, we took advantage of the apo-sGC activator drug, BAY 58-2667 (cinaciguat), which specifically binds to the empty heme binding pocket of apo-sGC and re-activates cGMP formation in an NO-independent manner ^42^. Indeed, up to 72 h exposure of PPAECs to DETA-NO (100 μM) increased apo-sGC activity, measured as BAY 58-2667-induced cGMP formation (Fig. 3C), and reduced sGC activity (Fig. 1D). To validate this mechanistic finding *in-vivo*, we re-examined the above already mentioned and utilized high-NO porcine ARDS model in which we had observed lower sGC protein levels and activity (see Fig. 1G, H). Consistent with our above *in-vitro* findings, apo-sGC activity was also increased in the high-NO porcine ARDS model (Fig. 3D) and established apo-sGC formation as one possible chronic mechanism of chronic NO in addition to total sGC protein loss (Fig. 3E).

## Discussion

Our findings close important gaps in our understanding of NO-cGMP signaling, in particular on the long-term effects of endogenously formed NO versus NO donors on sGC and the pathophysiological relevance of chronic NO for sGC regulation. We thus expand the previously observed notion that NO donors drugs can reduce sGC mRNA levels ^43^ to the protein level and importantly from pharmacology to endogenous NO. Previously, sGC protein levels were not consistently investigated or with antibodies of unclear specificity ^12,44^. Moreover, the functional consequences of PKG on cGMP levels were investigated only in some cases ^43,45^ or in relation to cGMP metabolism rather than its formation ^43,46,47^.

Surprisingly, not only pathological high levels of NO, as in our porcine ARDS model, but already low chronic endogenous NOS activity suppressed sGC protein and activity in an L-NAME reversible manner. These findings establish a previously not recognized delicate steady state in the interactions between NO and sGC, acutely stimulating and chronically limiting its expression and activity. On a positive note, under conditions of diminished NO synthesis, this may in turn rapidly upregulate sGC protein and activity as we have observed in the presence of the NOS inhibitor, L-NAME and *in-vivo* in eNOS^−/−^ mice. In this regard, previous data are controversial. For example, sGC activity was increased in eNOS^−/−^ mice ^13,48^ which agrees with our findings. However, other studies showed that neither sGC expression nor activity were changed in eNOS^−/−^ mice ^49,50^, or upon treatment with high dose NO donors ^51^. The reasons for this discrepancy are unclear.

Of therapeutic importance is also the previously not recognized risk of chronic use of NO donor drugs as they will lead to a downregulation of both sGC protein and activity. Together with the problematic pharmacokinetics and known, but entirely different phenomenon of pharmacokinetic tolerance due to lack of NO donor drug conversion to NO ^52–54^, this adds to the limitations of this widely used drug class. With the introduction of NO-independent sGC stimulators and cGMP elevating agents into clinical practice ^55^ there is now an alternative. Indeed, we show that the prototypic sGC stimulator and PDE inhibitor, YC-1, does not lead to sGC downregulation.

With respect to the underlying mechanisms, we initially considered two known mechanisms in NO-cGMP physiology, *i.e.* cGMP/PKG and thiol modification ^9,10^. Surprisingly, both could be excluded, which was reminiscent of an earlier observation where long-term exposure to an exogenous NO donor also reduced sGC activity in a manner that could not be recovered with thiol treatment ^14^. Instead, our findings suggest that endogenous and exogenous NO chronically induce a net shift from sGC to apo-sGC and that this is not only a pathophysiological mechanism but pertains to NO-cGMP physiology. This explains why apo-sGC activator-induced cGMP formation and functional effects are enhanced in but not exclusive to disease conditions ^7^. Nevertheless, the availability of sGC activator compounds allows now to overcome such conditions in which sGC protein and activity is diminished in favor of apo-sGC and still induce cGMP formation. As a limitation, other potential underlying mechanisms such as decreased mRNA ^43^ or increased degradation of sGC have not been addressed by our study ^56^ and cannot be excluded.

Our findings also add to our understanding of apo-sGC as a therapeutic target. Hitherto apo-sGC has been mainly studied by using the heme oxidant, ODQ, or by expressing enzyme where the proximal heme ligating histidine had been deleted ^57^. The mechanisms by which apo-sGC forms in pathophysiology were less clear. Now chronic exposure to (high) levels of NO can be considered one of these conditions. Whether this involves additional interactions with for example reactive oxygen species and from which source remains to be investigated. Certainly, intermediate compounds such as peroxynitrite would be candidate molecule to potentiate NO’s oxidative potential ^58^. Of note, the shift from sGC to apo-sGC is not 1-to-1. Some sGC appears to be lost due to inactivation beyond recovery by apo-sGC activators, e.g. by channeling into the ubiquitylation-proteasome pathway ^59^. Nevertheless, an apparent net shift from sGC to apo-sGC as main source of cGMP formation is a common denominator and has recently also been observed by us in another high NO model of ischemic stroke ^6^ and by others in an asthma model ^20^. In contrast to other observations, in our settings chronic NO incubation for 72 h unlike others for overnight ^20^, did have an effect on sGC β1 expression independent of S-nitrosylation.

In conclusion, our data suggest that both *in-vitro* and *in-vivo*, and both under physiological conditions and in disease NO self-limits its ability to induce cGMP formation via a chemical redox feedback which causes inactivation of sGC and an apparent net shift towards NO-insensitive apo-sGC. Our findings are of direct therapeutic importance as a pathological sGC/apo-sGC ratio can be treated with sGC activator compounds such as BAY58-2667 ^59^ thereby reinstalling cGMP synthesis and PKG signaling ^7,37^. Moreover, with respect to the long-established class of NO donor drugs and the use of inhaled NO a cautionary note needs to be added. Not only do they cause reversible tolerance, but, as we now find, also irreversible downregulation of sGC and apo-sGC formation. This explains the superiority of the novel, NO-independent sGC stimulators, at least in indications such as pulmonary hypertension ^15^.

## Materials and Methods

### Chemicals

Polyclonal antibodies specific for sGCα_1_ and sGCβ_1_ have been described elsewhere (30). Actin monoclonal antibody (Oncogene Research Products, Boston, USA); collagenase type CLS II (Merck, Netherlands); 8-bromo-cGMP (BIOLOG, Germany); L-NAME, DETA/NO, DEA/NO, IBMX and GTP (Enzo Life Sciences, Netherlands); BAY 58-2667 was synthesized as described ^60^. All other chemicals were of the highest purity grade available and obtained from Sigma or Merck (Netherlands). DETA/NO and DEA/NO were dissolved in 10 mM NaOH, BAY 58-2667 and YC-1 in DMSO.

### Tissue isolation

Tissues from i) 6- to 8-months old male PKG^−/−^ and age-matched control mice were obtained from Prof. Franz Hofmann, Department of Pharmacology and Toxicology at the Technical University Munich (genetic background 129/Sv) ^33^, and ii) 6- to 8-months old male eNOS^−/−^ mice and age-matched control were obtained from the Department of Physiology at Heinrich-Heine-Universität Düsseldorf (genetic background C57BL/6) ^36^. Animals’ care was in accordance with guidelines of Technical University Munich and Heinrich-Heine-Universität Düsseldorf. Experimental protocols were approved by the animal ethics committees of Technical University Munich and Heinrich-Heine-Universität Düsseldorf.

### Preparation of pulmonary arteries from a porcine ARDS model

Pulmonary arteries were removed immediately after death from an experimental porcine model of ARDS, as previously described ^35^. Pulmonary arteries were snap-frozen in liquid nitrogen and stored at minus 80°C or otherwise processed immediately to tissue powder and subsequently suspended in homogenization-buffer and homogenized in an Ultra Turrax at 4°C. These samples were then used further for protein determination, protein immune blots and sGC activity assays.

### PPAECs

Fresh porcine pulmonary arteries were obtained from a local slaughterhouse and maintained in phosphate-buffered saline (PBS; 10mM Na_2_HPO_4_, 1.8mM KH_2_PO_4_, 140mM NaCl, 2.7mM KCl, pH 7.4) at 37°C. PPAECs were isolated enzymatically by incubation of the aorta inner surface with collagenase type CLS II (0.5 mg/mL for 10 min at room temperature) and then collected in HEPES-buffered medium 199. After centrifugation (250 × g, 10 min) the pellet was re-suspended in growth medium (medium 199 supplemented with 10% fetal calf serum, 100 U/mL penicillin, 100 μg/mL streptomycin) and cells were propagated in coated plastic flasks and incubated (37°C, 6% CO_2_). Upon confluence, endothelial cell monolayers were sub-cultured in 35-mm (for Western blot) or 60-mm (for cGMP determination) gelatin coated dishes. Confluent cell monolayers from the second passage were used for experiments. The growth medium was replaced either every 12 or 24 hours if applicable containing the indicated compounds. After incubation time cells were subsequently used for sGC activity measurements or western blot analysis.

### Detection and quantification of sGC protein

Western blotting procedures were described previously ^61^. Briefly, cells were lysed in 250 μL Roti-Load sample buffer (ROTH, Karlsruhe, Germany), preheated to 95°C and then boiled for additional 10 min prior loading on SDS gel electrophoresis. Primary antibodies were diluted 1:4000 for anti-sGCα_1_ and 1:2000 for anti-sGCβ_1_ antibody in 3% dry milk in TBST and incubated with nitrocellulose membranes at 4°C over-night following challenge of membranes with secondary goat anti-rabbit antibody (1:2000 in 3% milk in TBST) conjugated to horseradish peroxidase (Dako A/S, Denmark). Immuno-complexes were visualized using an enhanced chemiluminescence kit (Amersham Pharmacia Biotech, Freiburg). Samples were quantified with a Kodak Imager Station 440 CF and with the NIH 1.6 software. All blots are standardized to ß-actin or GAPDH expression that was not affected by the treatments. Representative western blot examples are shown in Supplementary Fig. S3 online.

### Determination of sGC activity

To measure sGC activity, cells were stimulated with 250 μM DEA/NO or 10 μM BAY 58-2667 for 3 min at 37°C. Thereafter, cells were immediately lysed in 80 % ethanol. Cells were scraped and, after evaporation of ethanol, re-suspended in assay buffer and sonicated. Measurement of sGC activity in crude homogenates of porcine tissue was performed as previously described ^61^. Briefly, all samples were measured as the formation of cGMP at 37 °C during 10 min in a total incubation volume of 100 ml containing 50 mM triethanolamine-HCl (pH 7.4), 3 mM MgCl_2_, 3 mM glutathione, 1mM IBMX, 100mM zaprinast, 5 mM creatine phosphate, 0.25 mg/ml creatine kinase and 1mM or 0.5 mM GTP. The reaction was started by simultaneous addition of the sample and either DEA/NO or BAY 58-2667, respectively. After incubation of each sample for 10 min the reaction was stopped by boiling for 10 min at 95°C. Thereafter the amount of cGMP was subsequently determined by a commercial enzyme immunoassay kit (Enzo Life Sciences, Netherlands).

### Statistics

For comparisons students′ t-test or multiple comparisons one-way analysis of variance (ANOVA) was followed by Bonferroni’s test. Calculations were performed using GraphPad Prism 6.0 (GraphPad Software, San Diego, USA). All data are expressed as mean ± SEM. P-value < 0.05 was considered significant.

## Acknowledgements

We thank Prof. Hofmann for providing the PKG^−/−^ mice tissues; Helmut Müller, for expert technical assistance; Kirstin Wingler, Rob Hermans, Sabine Meurer and Merlijn Meens, for carefully reading and editing earlier versions of this manuscript.

## Author contributions

H.H.H.W.S. designed research; V.T.D., M.D., P.I.N., A.Gü., C.I.A performed research; A.Gü. and A.Gö. contributed new reagents/analytic tools; V.T.D., M.E., M.D., P.I.N., C.I.A. and H.H.H.W.S. analyzed data; and V.T.D., M.E., A.Gö. and H.H.H.W.S. wrote the paper.

## Data availability

All data needed to evaluate the conclusions in the paper are present in the paper or the Supplementary Materials.

