## Supplemental information for "A non-canonical chemical feedback self-limits nitric oxide-cyclic GMP signaling in health and disease"

### SUPPLEMENTARY FIGURES

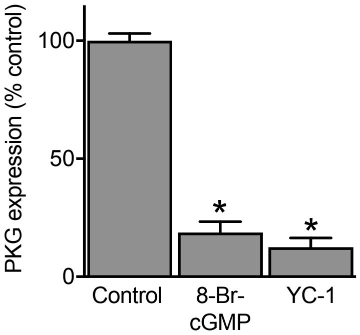


**Supplementary Fig. S1.** PPAECs display intact NO-cGMP-PKG signaling since treatment with 8-Br-cGMP (500 μM), or YC-1 for 72hrs reduced PKG expression (N=6-9). *P< 0.05 by one-way analysis of variance (ANOVA), data represent means ± SEM.

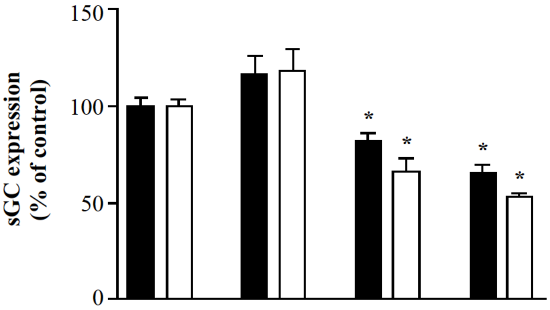


| | | | | |
|---|---|---|---|---|
| 100 μM DETA-NO | - | - | + | + |
| 10 μM Rp-8-Br-PET cGMPS | - | + | - | + |

**Supplementary Fig. S2.** Inhibition of PKG does not prevent DETA-NO induced downregulation of sGC$\alpha_1$ and sGC $\beta_1$. Treatment of PPAECs for 72h with either 100 μM DETA-NO, or with the PKG inhibitor Rp-8-Br-PET-cGMPS (10 μM), or a combination thereof resulted in down regulation of sGC$\alpha_1$ (solid bars) and sGC$\beta_1$ (open bars) level that could not be reversed by Rp-8-Br-PET-cGMPS. *P< 0.05 by one-way analysis of variance (ANOVA), data represent means ± SEM

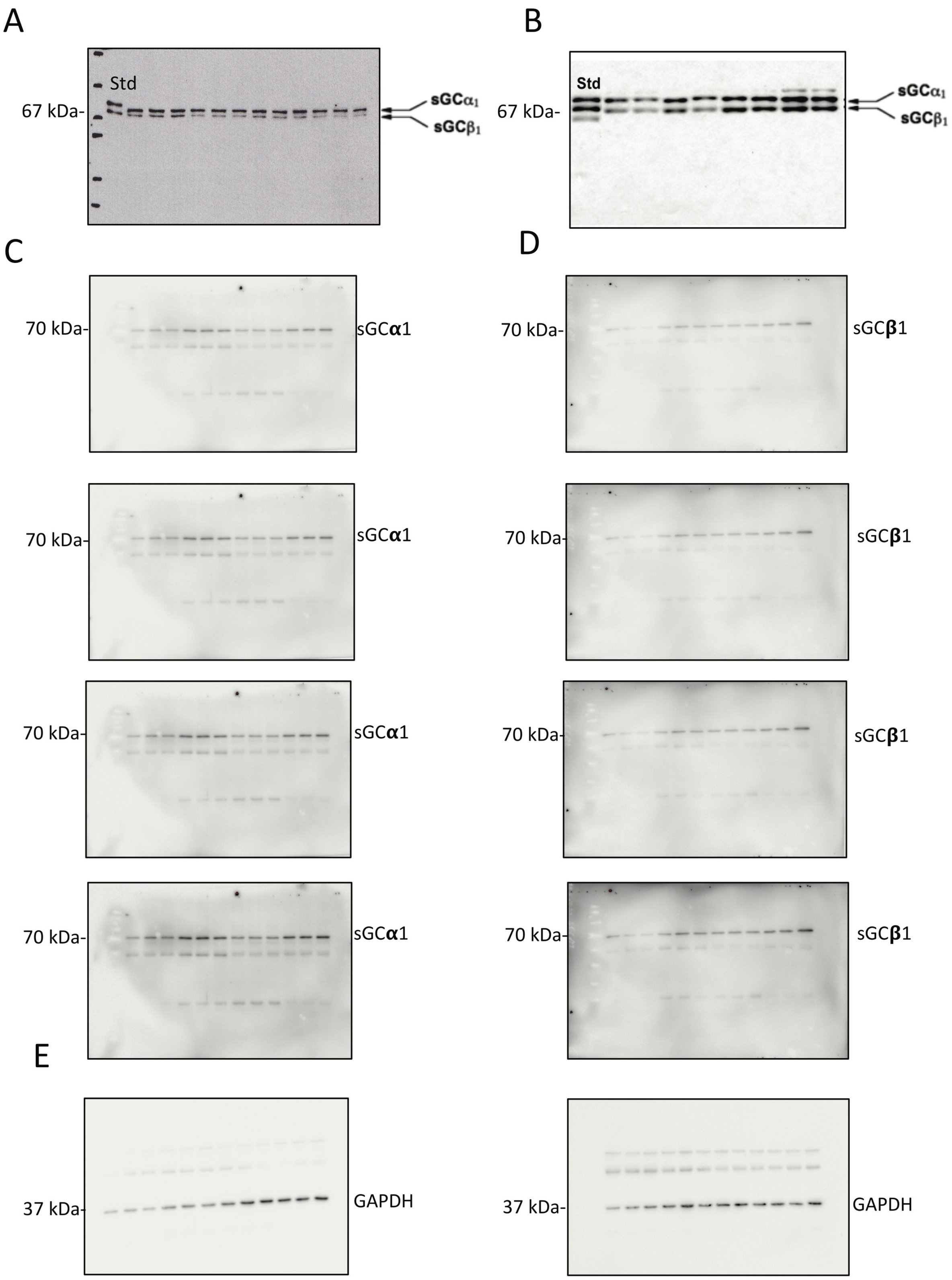


**Supplementary Fig. S3.** Representative Immune detection blots of controls, sGC$\alpha_1$ and sGC$\beta_1$. A. PPAECs. B. Mouse. C-E. ARDS (Multiple exposures).
